# CMTM6 suppresses cell-surface expression of death receptor FAS in mice but not in humans

**DOI:** 10.1101/2025.05.12.653457

**Authors:** Tereza Semberova, Michaela Pribikova, Helena Kissiova, Tijana Trivic, Ondrej Stepanek, Peter Draber

**Affiliations:** Laboratory of Immunity & Cell Communication, Division BIOCEV, First Faculty of Medicine, Charles University, Vestec, Czech Republic; Department of Immunobiology, University of Lausanne, Epalinges, Switzerland; Laboratory of Adaptive Immunity, Institute of Molecular Genetics of the Czech Academy of Sciences, Prague, Czech Republic

## Abstract

The transmembrane protein CMTM6 was shown to promote plasma membrane expression of immune checkpoint PD-L1, an important suppressor of anti-tumor immunity. Targeting of CMTM6 was proposed as a strategy to decrease surface PD-L1 and trigger cytotoxicity against tumors. In accord, ablation of CMTM6 in mouse cancer models was shown to efficiently suppress tumor growth in a manner partially independent of PD-L1, which suggested that CMTM6 might regulate other proteins involved in anti-tumor immunity. Using mass spectrometry, we discovered that mouse CMTM6 strongly associates with the cell death receptor FAS and negatively regulates its expression in mice. Deletion of CMTM6 promotes FAS membrane localization and renders murine cells sensitive to FASL-mediated cytotoxicity. However, the interaction between CMTM6 and FAS is lost in human cells due to the difference in three amino acids at the boundary of the FAS extracellular and transmembrane domains. Altogether, our data urge caution when transferring promising data regarding the targeting of CMTM6 from mouse cancer models to potential human therapies.

## Introduction

Induction of programmed cell death is critical to maintain immune homeostasis and to remove damaged or infected cells. FAS is a widely expressed transmembrane receptor that can trigger apoptotic cell death upon binding of FAS ligand (FASL). In contrast, FASL is expressed mainly by activated T cells and NK cells (Nagata, 1997). FAS and FASL are important in controlling lymphocyte numbers. The deficiency in FAS or FASL proteins causes autoimmune lymphoproliferative syndrome (ALPS) in humans (Rieux-Laucat *et al*, 2018). In accord, ablation of either protein triggers severe lymphoproliferative disease in mice (Adachi *et al*, 1995; Karray *et al*, 2004; Takahashi *et al*, 1994; Watanabe-Fukunaga *et al*, 1992). Apart from regulating immune homeostasis, FAS-induced apoptosis is also employed by the immune system to kill virus-infected or transformed cells. Upon stimulation, the FAS intracellular death domain assembles a death-inducing signaling complex (DISC) containing adaptor FADD that recruits caspase-8, caspase-10, and their regulator cFLIP. Autoproteolytic cleavage of caspase-8/-10 releases their active forms into the cytoplasm, thus initiating the caspase cascade, and leading to apoptosis (Peter *et al*, 2015).

FAS stimulation can also trigger non-apoptotic signaling, especially activation of NF-κB and production of proinflammatory cytokines (Cullen *et al*, 2013; Davidovich *et al*, 2023; Siegmund *et al*, 2017). While immune cells can use FASL to kill tumor cells, some cancers are highly resistant to FAS-induced cell death and instead employ FAS-induced signaling to enhance their growth and invasiveness (Barnhart *et al*, 2004). Although the precise mechanism guiding the outcome of FAS stimulation is still under investigation, internalization of FAS appears crucial for the induction of cell death (Lee *et al*, 2006; Magri *et al*, 2024). Given the ability of FAS to trigger either cell death or activation and migration of cells, both FAS agonists and antagonists are being tested as therapeutic strategies to suppress tumor growth and modulate inflammation (Risso *et al*, 2022).

Immune surveillance of tumors involves T cells that can recognize and destroy cancer cells. However, their activity within the tumor microenvironment is often suppressed by the immune checkpoint protein programmed death 1 (PD-1), which binds to its ligand PD-L1, commonly expressed on cancer cells. Stimulation of PD-1 suppresses T cell activity, leading to their exhaustion (Morad *et al*, 2021). The surface expression of PD-L1 is regulated by association with small four-membrane domain-containing protein CMTM6 and, to a lesser degree, its homologue CMTM4 (Burr *et al*, 2017; Dai *et al*, 2021; Mezzadra *et al*, 2017). Indeed, targeting CMTM6 in several mouse cancer models led to enhanced anti-tumor cytotoxicity, which was only partially caused by decreased PD-L1 expression, since blocking PD-L1 together with CMTM6 depletion had a substantially stronger effect than each treatment alone (Long *et al*, 2023). This indicated that in addition to enhancing PD-L1, CMTM6 protects mouse cancer cells by regulating the expression of other protein(s).

Here, we aimed to identify the interacting partners of mouse CMTM6 compared to other CMTM family members to search for other immunologically relevant proteins they might regulate. We showed that endogenous CMTM6 strongly binds to mouse FAS and suppresses its plasma membrane localization, therefore reducing sensitivity to FASL-driven apoptosis. However, human FAS does not bind CMTM6 due to a difference in three amino acids in the transmembrane and extracellular region of human and mouse FAS. Altogether, our data demonstrate different regulation of FAS membrane expression between mouse and human cells and suggest that targeting human CMTM6 might not have the same benefit as targeting the protein in mouse tumor models.

## Results

### Mouse FAS is strongly associated with CMTM6

CMTM4 and CMTM6 proteins were previously reported as important regulators of immune responses and anti-tumor immunity due to directly binding and regulating plasma membrane expression of several proteins, such as PD-L1 (Burr *et al*., 2017; Dai *et al*., 2021; Mezzadra *et al*., 2017), IL-17 receptor C (IL-17RC) (Knizkova *et al*, 2022; Ni *et al*, 2024), or the epidermal growth factor receptor (EGFR) (Xu *et al*, 2025). In here, we employed mass spectrometry to elucidate whether CMTM4 and CMTM6 bind other immunologically relevant proteins that might play a role in regulating tumor immunosurveillance.

We first prepared retroviral vector coding for CMTM4 fused at the C-terminus to the 2xStrep 3xFlag (SF) tag. Expression of CMTM4-SF in *Cmtm4*^KO^ mouse stromal ST2 cells fully rescued the expression of surface IL-17RC, indicating that the addition of the tag does not interfere with the CMTM4 function (Fig. S1A). Next, we prepared six ST2 cell lines expressing CMTM3 to CMTM8 fused to the SF-tag. We excluded CMTM1 and CMTM2 from our study because these proteins are testis-specific according to the Human Protein Atlas (Uhlen *et al*, 2015) and unstable upon overexpression in ST2 cell lines (Knizkova *et al*., 2022). We purified individual CMTM family members using tandem affinity purification and analyzed associated proteins by mass spectrometry (Table S1). Protein quantification was based on Top3 intensity, which sums the three most intense peptides derived from each protein. We identified PDL1 among the top 20 interactors of CMTM6 (Fig. 1A). Similarly, IL-17RC was a very strong interactor of CMTM4 (Fig. S1B), which validated our experimental approach. Some members of the CMTM family interacted together, such as CMTM4 and CMTM6, suggesting they might create membrane nanodomains, as was observed for proteins of the tetraspanin family (van Deventer *et al*, 2021) (Fig. S1C). Importantly, we observed that cell-death receptor FAS strongly interacted with CMTM6 and was very weakly associated with CMTM4 but not with other CMTM family members (Fig. 1A-B). Finally, we confirmed the strong interaction between CMTM6-SF and murine FAS via immunoblotting (Fig. 1C).

**Figure 1.**
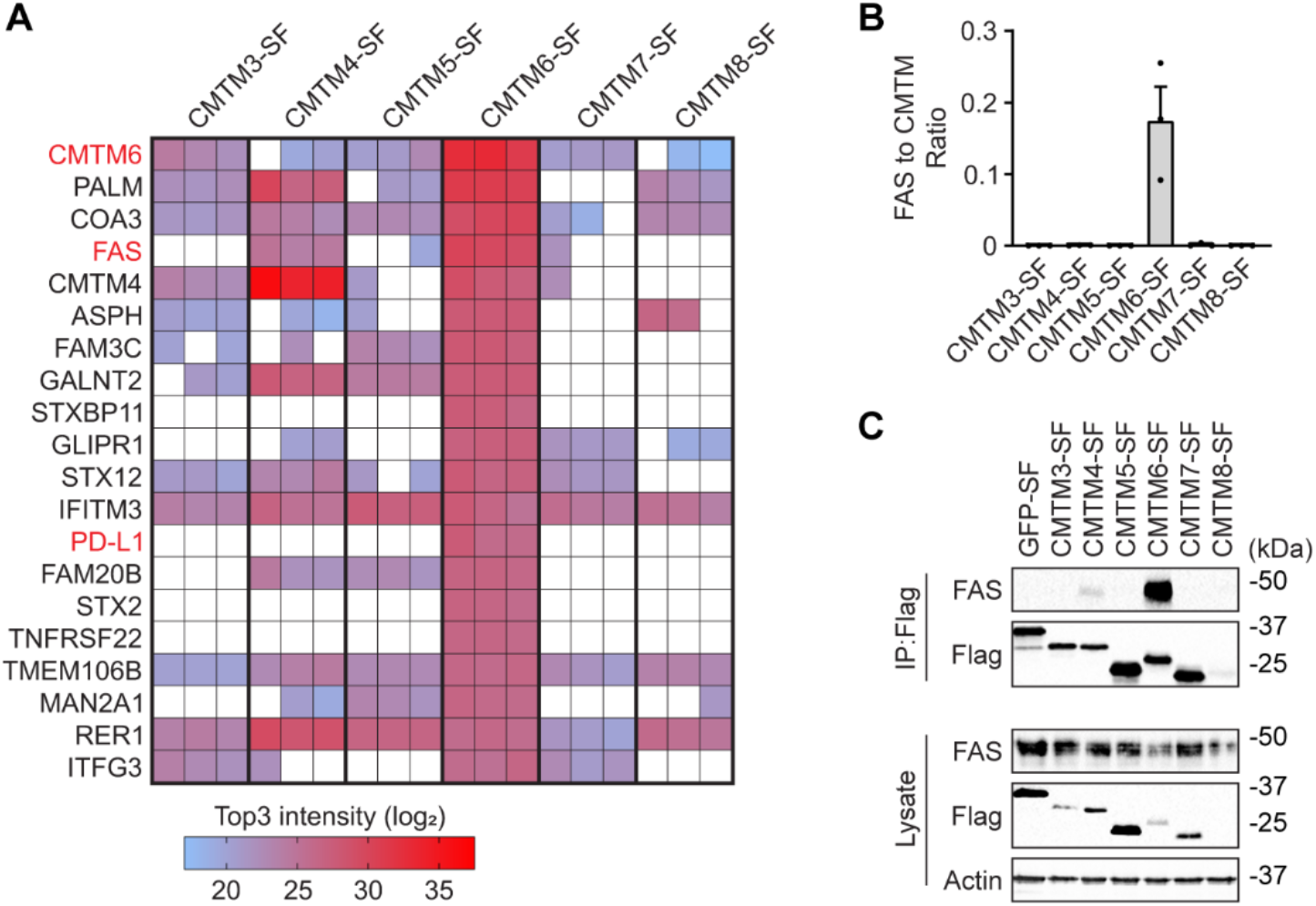
CMTM6 associates with FAS in mouse cells. **A**. Mass spectrometry analysis of the indicated Strep-Flag (SF)-tagged CMTM family members isolated from mouse ST2 cells via tandem affinity purification. The heatmap shows the most abundant CMTM6 interactors that were not detected in control GFP-SF samples, based on Top3 intensities from 3 independent experiments. **B**. Quantification of the Top3 intensity ratio of FAS relative to individual SF-tagged CMTM family members. **C**. Samples isolated via Flag immunoprecipitation from ST2 cells expressing the indicated SF-tagged CMTM family members or GFP were analyzed by immunoblotting. Data are representative of 3 independent experiments.

To validate our data, we expressed murine FAS fused to SF-tag at the C-terminus in ST2 cells and analyzed the FAS-interactome via mass spectrometry upon tandem affinity purification. We observed that in unstimulated cells, CMTM6 was the major interactor of FAS (Table S2 and Fig. 2A). We validated their interaction also using immunoblotting (Fig. 2B). We also detected very weak binding of CMTM4 (Fig. 2B). CMTM6 and CMTM4 have four transmembrane domains with short extracellular or cytoplasmic sequences, which indicated that they might bind to transmembrane or juxtamembrane domain of murine FAS. To test this hypothesis, we expressed a chimeric protein in which the FAS transmembrane and juxtamembrane parts were exchanged for the corresponding sequence from cell death receptor TRAILR2 that does not interact with CMTM family members (Fig. 2C and S1D). This led to the complete loss of CMTM6 and CMTM4 binding (Fig. 2D). In contrast, TRAILR2 harboring the transmembrane and juxtamembrane domains of murine FAS strongly interacted with CMTM6, and we detected weak binding of CMTM4 (Fig. 2C-D). Altogether, these data showed that FAS interacted with both CMTM4 and CMTM6 via the transmembrane region or potentially via several amino acids flanking this domain.

**Figure 2.**
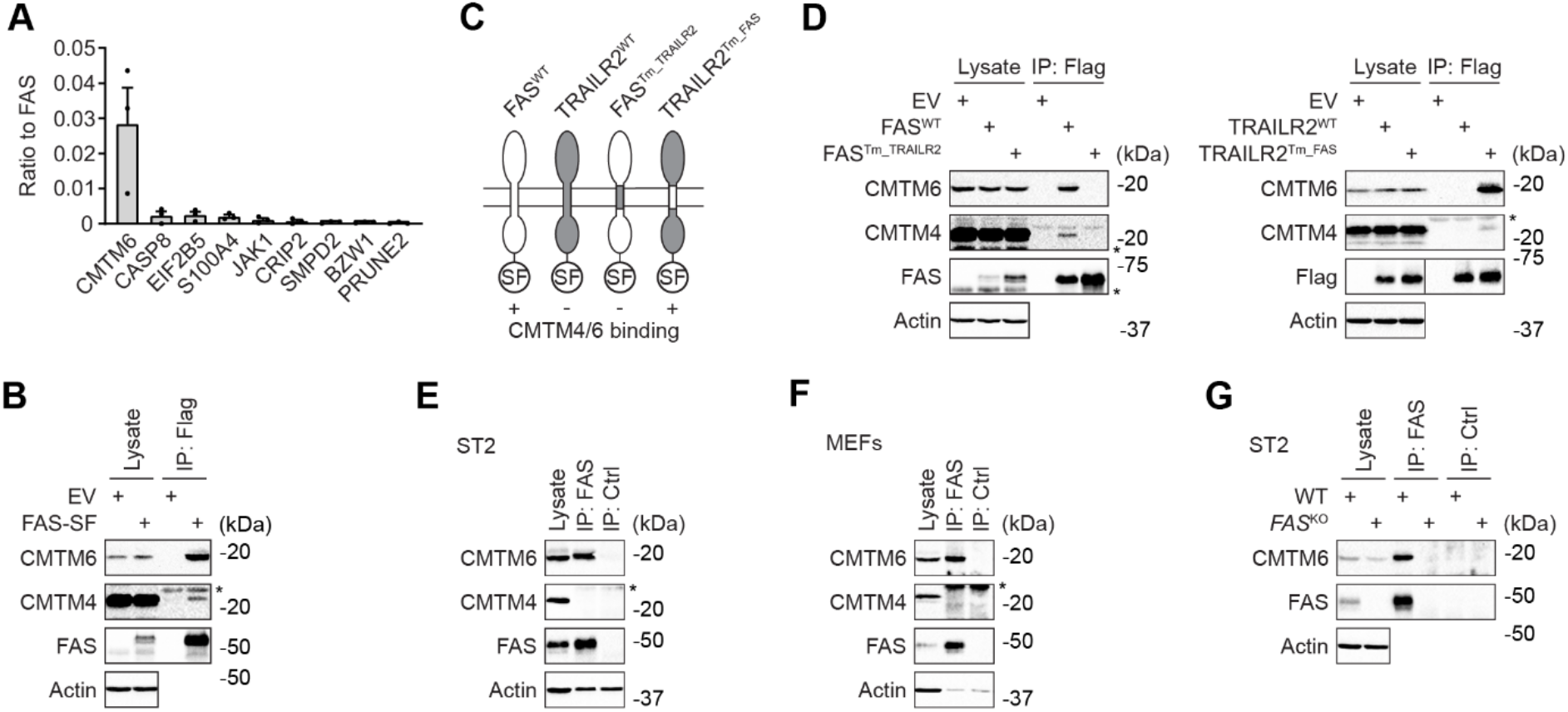
Mouse FAS binds strongly to CMTM6 via the transmembrane domain. **A**.Mass spectrometry analysis of the SF-tagged mouse FAS isolated from mouse ST2 cells via tandem affinity purification. The most abundant FAS interactors not detected in control GFP-SF samples are shown, based on Top3 intensity from 3 independent experiments. **B**. ST2 cells expressing SF-tagged mouse FAS or empty vector (EV) were lysed, subjected to Flag immunoprecipitation, and analyzed by immunoblotting. **C**. Schematic representation of mouse FAS and TRAILR2 chimeric proteins expressed in ST2 cells. **D**. SF-tagged FAS, TRAILR2, and the indicated chimeric proteins were purified via Flag immunoprecipitation and analyzed by immunoblotting. **E-G**. Lysates from the indicated cell lines were subjected to immunoprecipitation with anti-FAS or isotype control antibodies and analyzed by immunoblotting. Data are representative of two (E-G), three (A, B), and five (D) independent experiments. *, nonspecific band.

The immunoprecipitation of endogenous FAS showed a strong interaction with CMTM6, but we did not detect CMTM4 in mouse ST2 cells or mouse embryonic fibroblasts (MEFs), indicating that CMTM6 is the major interacting partner of endogenous protein (Fig. 2E-F). Finally, we co-immunoprecipitated CMTM6 using anti-FAS antibody from ST2 wild-type (WT) cells but not from FAS-deficient cells (Fig. 2G), which further confirmed the specificity of the interaction. Combined, these data established a strong and specific interaction of endogenous FAS with CMTM6 in mouse cells.

### CMTM6 suppresses FAS membrane localization

CMTM6 was previously shown to promote PD-L1 plasma membrane localization. FAS is an important regulator of immune homeostasis, and FAS-deficient mice develop lymphoproliferation (Peter *et al*., 2015). To evaluate whether CMTM6 is important in regulating immune system homeostasis, we analyzed the immune system in *Cmtm6*^−/−^ mice (gating strategy is shown in Fig. S2). However, we did not detect any changes in immune cell homeostasis. There were no differences in spleen weight or cellularity between WT and *Cmtm6*^−/−^ mice (Fig. 3A, S3A). The proportion of splenic CD3^+^ T cells and the ratio between cytotoxic CD8^+^ T cells, helper CD4^+^ T cells, and regulatory T cells (Tregs) was unchanged (Fig. 3B). We did not detect an increased proportion of splenic memory CD4^+^ T cells in *Cmtm6*^−/−^ mice (Fig. 3C). Similarly, we observed no changes in the proportion of memory splenic CD8^+^ T cells and antigen inexperienced memory-like CD8^+^ T cells (AIMT) that did not encounter antigen (Moudra *et al*, 2021) (Fig. 3D). We observed the same results in lymph nodes (Fig S3A-D), indicating that CMTM6 is not required to regulate the general T-cell homeostasis.

**Figure 3.**
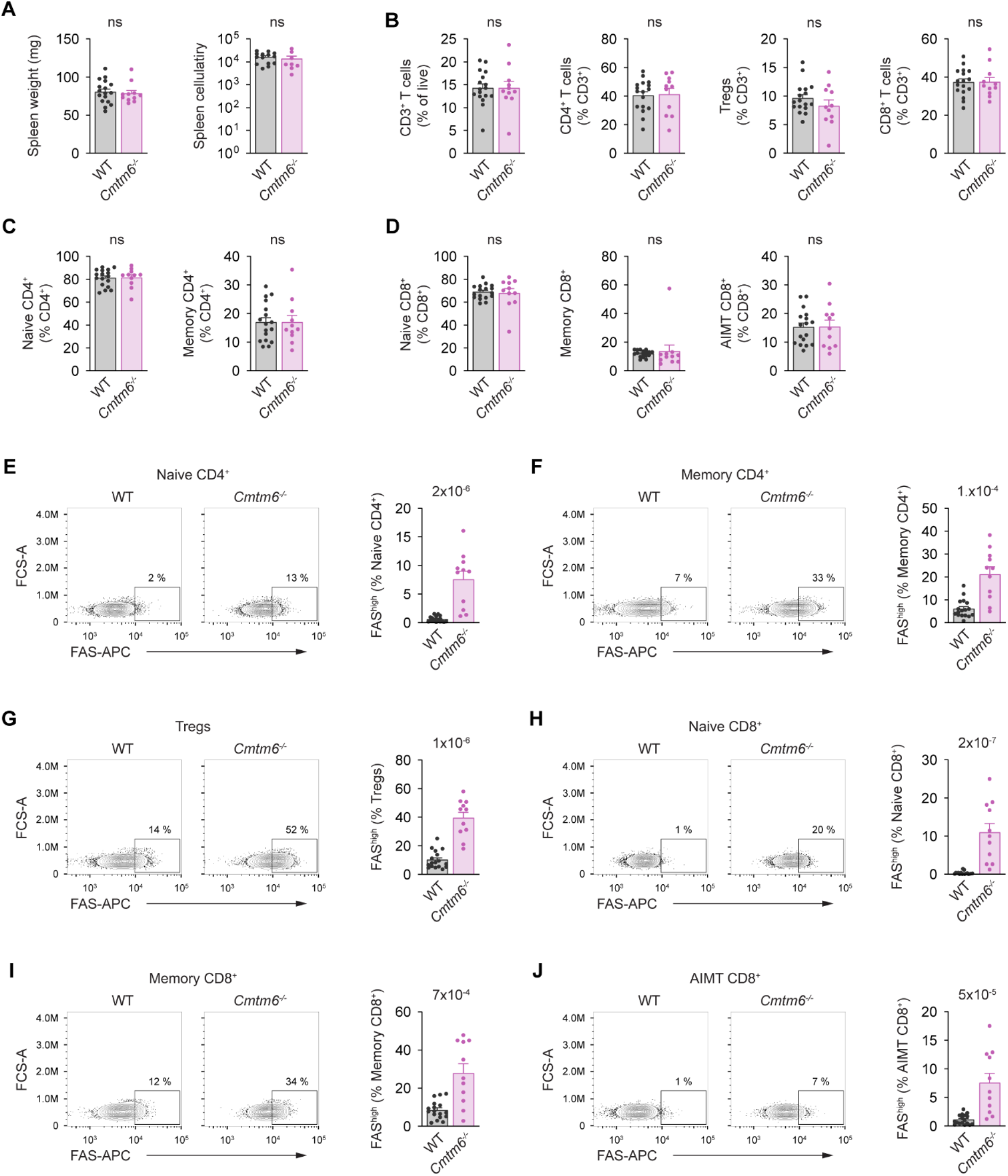
CMTM6 suppresses FAS surface levels in mouse splenic T cells. **A**.Weight and cellularity of spleens isolated from 8-12-week-old *WT* and *Cmtm6*^*-/-*^ mice. **B**. Flow cytometry analysis of splenic T cells from 8-12-weeks-old *WT* and *Cmtm6*^*-/-*^ mice. T cells (gated as CD3^+^) were subdivided into conventional CD4^+^ (CD4^+^, FOXP3^−^), Tregs (CD4^+^, FOXP3^+^), and conventional CD8^+^ (CD8^+^). **C**. Conventional CD4^+^ T cells were further gated as naïve (CD4^+^, FOXP3^−^, CD44^−^, CD62L^+^) and memory (CD4^+^, FOXP3^−^, CD44^+^, CD62L^−^). **D**. Conventional CD8^+^ T cells were gated as naïve (CD8^+^, CD44^−^), memory (CD8^+^, CD44^+^, CD49d^+^), and antigen-inexperienced memory T (AIMT) (CD8^+^, CD44^+^, CD49d^−^) cells. **E-J**. Surface FAS levels in the indicated subsets of splenic T cells. Data are presented as mean + SEM, n=8 mice per group. Two-tailed Mann-Whitney test. ns, not significant.

The analysis of T cells showed that Cmtm6^−/−^ T cells have markedly elevated expression of surface FAS compared to WT cells. We observed highly increased surface expression of FAS in both naïve and memory CD4^+^ T cells (Fig. 3E-F), Tregs (Fig. 3G), and in naïve, memory, and AIMT CD8^+^ T cells (Fig. 3H-J) isolated from the spleen. Similarly, increased FAS membrane expression was observed in T cells isolated from Cmtm6^−/−^ lymph nodes (Fig. S3E-G). These results show that CMTM6 binding to FAS suppresses its membrane expression without affecting immune homeostasis. This was intriguing, as CMTM6 has an opposite effect on the immunosuppressive protein PD-L1 and promotes its membrane localization (Burr *et al*., 2017; Mezzadra *et al*., 2017).

### CMTM6 suppresses FAS-induced cell death in mouse cells

To validate that CMTM6 suppressed FAS expression on the cell surface, we isolated MEFs from WT or *Cmtm6*^−/−^ littermates. Indeed, the surface FAS was substantially increased in *Cmtm6*^−/−^ cells (Fig. 4A). Stimulation of *Cmtm6*^−/−^ MEFs with hexameric FAS ligand (Hex-FASL) led to markedly enhanced induction of apoptosis compared to WT controls (Fig. 4B). In accord, we detected increased cleavage of caspase-8 and caspase-3 in *Cmtm6*^−/−^ cells upon FASL stimulation compared to WT cells (Fig. 4C). As expected, reconstitution of *Cmtm6*^−/−^ MEFs with CMTM6 led to decreased FAS expression compared to cells reconstituted with empty vector (EV) only (Fig. 4D). In accord, enhanced induction of cell death upon FASL stimulation in *Cmtm6*^−/−^ MEFs and activation of caspase-8 and caspase-3 were rescued upon reconstitution of cells with CMTM6, but not EV (Fig. 4E-F). Altogether, our data demonstrate that CMTM6 suppresses FAS expression on the surface of mouse cells, which correlates with reduced FASL-induced activation of apoptosis.

**Figure 4.**
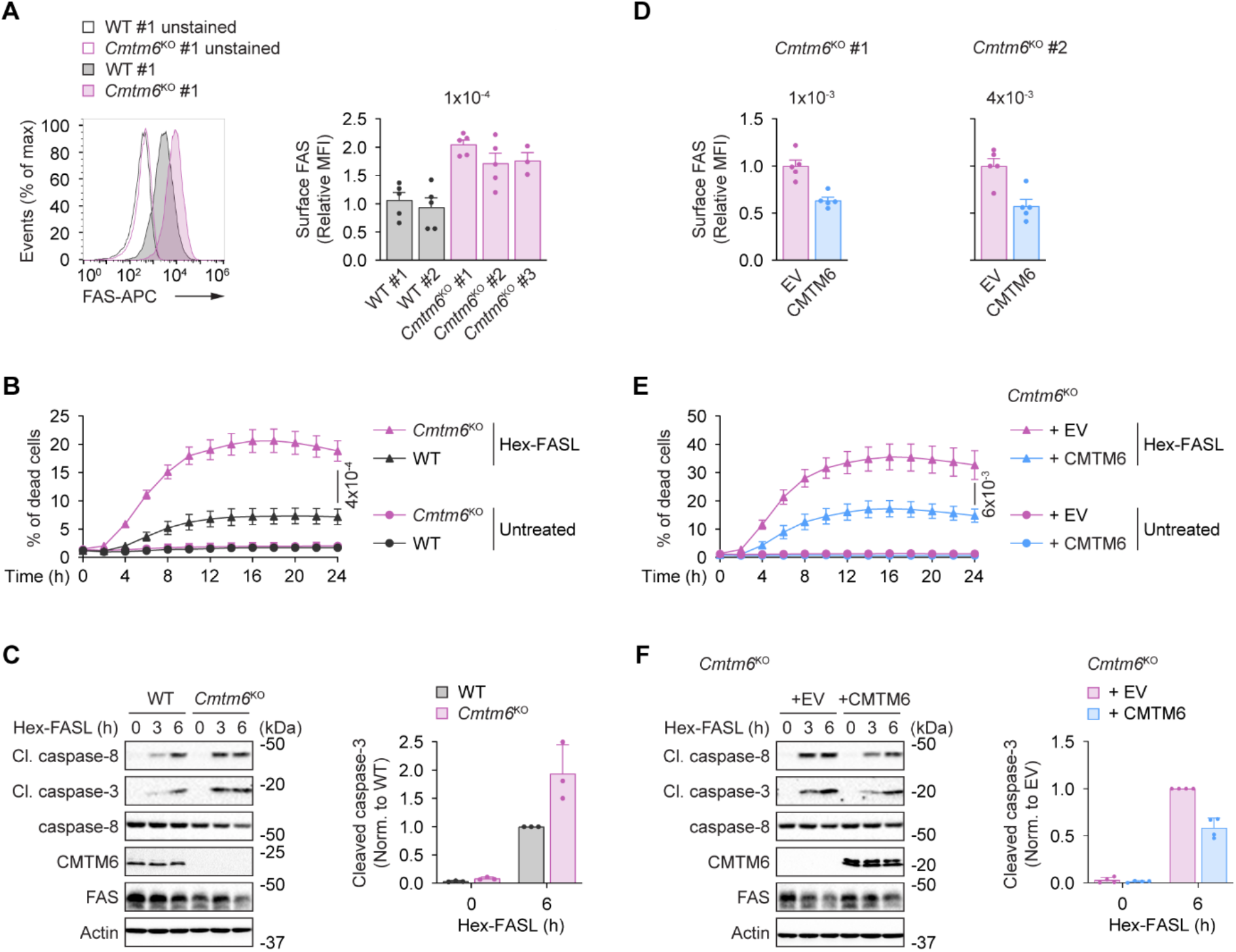
CMTM6 suppresses FAS-induced cell death. **A**.Flow cytometry analysis of surface FAS expression in MEFs isolated from *WT* and *Cmtm6*^*-/-*^ sibling embryos. **B**. Cell death induction in WT or *Cmtm6*^*-/-*^ MEFs (clones #1 and #2) stimulated or not with Hex-FASL (500 ng/μl). Cell death was monitored every 2 hours using Incucyte. **C**. Immunoblot analysis of lysates from WT or *Cmtm6*^*-/-*^ MEFs stimulated with Hex-FASL (500 ng/μl) for the indicated time points. **D**. Flow cytometry analysis of surface FAS expression in *Cmtm6*^*-/-*^ MEFs transduced with either an empty vector (E.V.) or a vector expressing untagged murine CMTM6. **E**. Cell death induction in *Cmtm6*^*-/-*^ MEFs (clones #1 and #2) reconstituted or not with CMTM6 that were stimulated or not with Hex-FASL (500 ng/μl). Cell death was monitored every 2 hours using Incucyte. **F**. Immunoblot analysis of lysates from *Cmtm6*^*-/-*^ MEFs reconstituted or not with CMTM6 that were stimulated with Hex-FASL (500 ng/μl) for the indicated time points. Data are representative of three (C), four (F), five (A, D), and six (B, E) independent experiments. One-way ANOVA (A) and an unpaired two-tailed t-test (B,D,E) statistical test. MFI, median fluorescent intensity.

### CMTM6 does not interact with human FAS

Our data indicated that CMTM6 has an important role in regulating FAS in mouse cells, and we aimed to validate our findings in human cells. To our surprise, we detected no interaction between endogenous human FAS and CMTM6 in HeLa cells (Fig. 5A) or in HEK293 cells (Fig. 5B). To exclude that technical issues had caused the lack of interaction between the two proteins in human cells, we expressed human FAS or PD-L1 fused to SF-tag in HeLa cells. Immunoprecipitation of cell lysates using anti-Flag antibody revealed strong interaction of CMTM6 with overexpressed human PD-L1 but not with human FAS (Fig. 5C). Altogether, these data indicated that FAS and CMTM6 do not interact in human cells. Mouse and human CMTM6 are very similar, sharing 83% of identical amino acids (Fig. S4A). This suggests that the differences between human and mouse FAS regarding CMTM6 binding are likely due to the differences in amino acid composition of FAS transmembrane and juxtamembrane domains.

**Figure 5.**
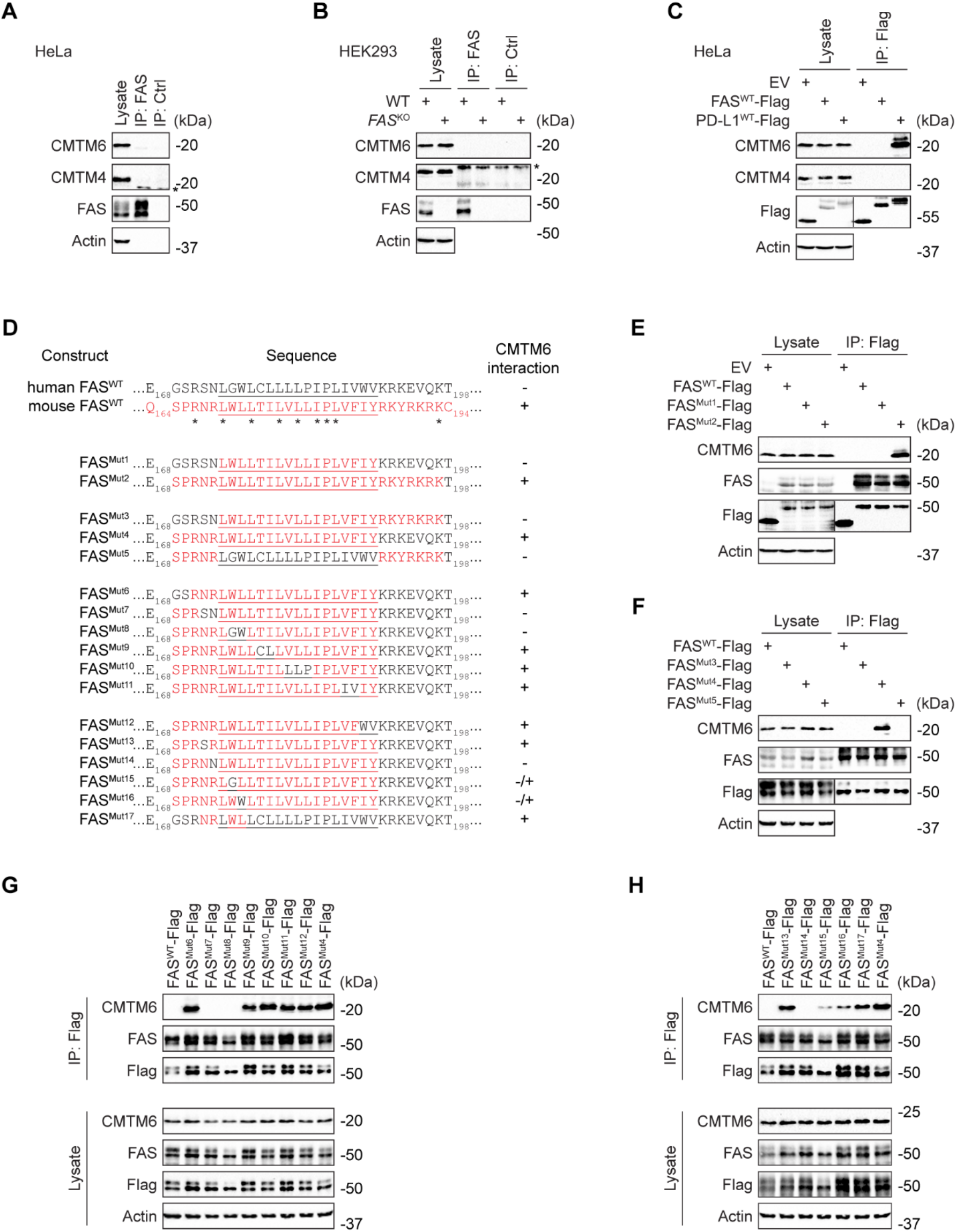
Human FAS does not interact with CMTM6. **A-B**. Lysates from the indicated cell lines were subjected to immunoprecipitation using anti-FAS or isotype control antibodies and analyzed by immunoblotting. **C**. HeLa cells expressing SF-tagged human FAS, human PD-L1, or empty vector (EV) were lysed, subjected to Flag immunoprecipitation, and analyzed by immunoblotting. **D**. Sequences of swap mutants between human and mouse FAS. The transmembrane domain is underlined. **E-H**. Lysates from HeLa cells expressing the indicated SF-tagged FAS mutants were subjected to Flag immunoprecipitation and analyzed by immunoblotting. Data are representative of two (A-B, E-H) and three (C) independent experiments.

We performed a series of experiments in which we swapped parts of human FAS for corresponding sequences in mouse FAS to identify what amino acid changes are causing the loss of CMTM6 binding to human FAS (Fig. 5D). First, we noted that human FAS^Mut1^, in which we exchanged only the transmembrane domain with its mouse counterpart, was insufficient to bind CMTM4. In contrast, human FAS^Mut2,^ in which we exchanged the transmembrane domain and the surrounding regions with mouse FAS, interacted strongly with CMTM6 in human HeLa cells (Fig. 5E). Subsequently, we demonstrated that human FAS^Mut4^, in which we swapped only the transmembrane and proximal extracellular part with the mouse counterpart, enabled CMTM6 binding (Fig. 5F).

Next, we started changing amino acids in the mouse FAS transmembrane and proximal extracellular part to those found in human FAS. The FAS^Mut7-8^, which harbored mutations of four amino acids at the boundary of the extracellular and transmembrane parts, lost the ability to bind to CMTM6 (Fig. 5G). Finally, detailed mapping in FAS^Mut14-16^ showed that three amino acids in mouse FAS that differ from those in humans are required for the interaction (Fig. 5H). Importantly, human FAS^Mut17^ with four amino acid changes to mouse homologues, i.e., S172N, N173R, G175W, and W176L, fully regained the ability to bind CMTM6 (Fig. 5H). These results show that the extracellular and transmembrane domain boundary of mouse FAS is required for interaction with CMTM6. Since this sequence is different in human FAS, the interaction is lost.

Finally, to analyze the interactome of human CMTM6, we retrovirally expressed SF-tagged human CMTM6 in HeLa cells, isolated CMTM6-SF via tandem affinity purification, and analyzed the associated proteins via mass spectrometry (Table S3). We observed that the strongest interactor of human CMTM6 was human immunoregulatory protein CD58 (Fig. S4B). We did not detect PD-L1 in our samples, as its expression is only induced upon IFNγ stimulation in HeLa cells (Fig. S4C). CD58 is an important activator of anti-tumor immunity, which is missing in mouse cells and was previously shown to interact with human CMTM6 (Ho *et al*, 2023; Miao *et al*, 2023). The association with human FAS was not detected. In accord, while CMTM6-deficient HeLa cells had decreased surface expression of CD58 and also decreased expression of PD-L1 in IFNγ-stimulated cells, the expression of FAS was not changed (Fig. S4D). Altogether, our data established that CMTM6 regulates mouse, but not human, FAS expression.

## Discussion

In this work, we analyzed the interactome of six members of the mouse CMTM family, CMTM3-8. We did not include mouse CMTM1, CMTM2a, and CMTM2b as these proteins are unstable upon overexpression in ST2 cells (Knizkova *et al*., 2022). We also noticed that individual members of the CMTM family interacted with each other. This suggests that CMTM members may associate in the membrane, similar to the proteins of the tetraspanin family, which also contain four transmembrane domains and short intracellular sequences, but differ in the presence of a large ectodomain. Tetraspanins create so-called tetraspanin nanodomains, which are important for the organization of membrane proteins and thus regulating diverse functions such as cell migration, cell signaling, or protein trafficking (Susa *et al*, 2024). However, whether the members of the CMTM family create functional nanodomains remains to be established.

Our major focus was the identification of proteins that interact with mouse CMTM6 and could have a role in regulating tumor growth. We detected a relatively limited number of immunologically important transmembrane receptors. Most interestingly, we identified FAS as a strong interactor of mouse CMTM6, and we also weakly detected FAS in the mouse CMTM4 interactome, while it did not associate with other CMTM members. Our data indicated that CMTM6 binds to the transmembrane and juxtamembrane domain of FAS. PD-L1 employs a similar interaction mode to bind CMTM6 (Mezzadra *et al*., 2017), and therefore, it is possible that FAS competes with PD-L1 for CMTM6 binding.

The *Cmtm6*^−/−^ mice analyzed in this work showed no detectable defect in T-cell homeostasis, indicating that CMTM6 is dispensable for T cell development. CMTM4 and/or CMTM6 were shown to promote membrane localization of several proteins, such as PD-L1, CD58, IL-17RC, or EGFR (Burr *et al*., 2017; Knizkova *et al*., 2022; Mezzadra *et al*., 2017; Xu *et al*., 2025). Therefore, our initial assumption was that CMTM6 would also enhance FAS membrane localization. To our surprise, this was not the case, as T cells isolated from *Cmtm6*^−/−^ mice had markedly enhanced membrane localization of FAS. In accord, *Cmtm6*^−/−^ mice did not suffer from the lymphoproliferative disease that is typical for animals with impaired FAS expression or function (Watanabe-Fukunaga *et al*., 1992). Interestingly, CMTM4 was shown to promote endocytosis of VE-cadherin and its internalization (Chrifi *et al*, 2019). It is possible that CMTM6 similarly facilitates the removal of FAS from the plasma membrane in mice, and therefore, its deletion may lead to FAS accumulation on the cell surface. The role of CMTM6 as a negative regulator of FAS surface expression is further supported by the increased sensitivity of CMTM6-deficient mouse cells to FASL-induced cell death. However, in human cells, FAS is not associated with CMTM6 due to the changes in three amino acids at the extracellular and transmembrane domain boundary. For this reason, we decided not to further study the mechanism of mouse FAS regulation by CMTM6.

Mice are widely used animals to study tumor development. Since CMTM6 promotes PD-L1 expression, the role of CMTM6 in tumor progression was extensively studied. Ablation of CMTM6 decreased tumor growth in various mouse models due to increased T-cell cytotoxicity (Long *et al*., 2023). This effect was correlated with the decreased expression of immunosuppressive immune checkpoint PD-L1. Still, other mechanisms contributed to the increased cytotoxicity against CMTM6-negative tumors, as the combined deficiency of CMTM6 and PD-L1 led to a stronger anti-tumor immune reaction than the absence of either protein alone (Long *et al*., 2023). It is possible that CMTM6-deficient mouse tumor cells are prone to T-cell-mediated cytotoxicity due to enhanced plasma membrane expression of FAS. In support of this hypothesis, tumor cells can decrease FAS surface expression via enhanced endocytosis and lysosomal degradation. Slowing FAS endocytosis or promoting its recycling to the plasma membrane led to increased FAS surface expression and enhanced FASL-induced killing of target cells (Kural *et al*, 2024; Sharma *et al*, 2019). The dual role of CMTM6 in enhancing expression of immunosuppressive PD-L1 while suppressing expression of death-inducing FAS in mouse cells is an intriguing mechanism by which CMTM6 might prevent cytotoxic attack against self and might therefore play an important role in preventing anti-tumor immunity in mice.

In contrast, human FAS is not regulated by CMTM6. Instead, we noted strong interaction between human CMTM6 is a costimulatory receptor CD58. This is in accord with previous reports showing that in human cells, CMTM6 is required for membrane expression of both PD-L1 and CD58 (Ho *et al*., 2023; Miao *et al*., 2023). CD58 is an adhesion molecule whose ligand, CD2, is expressed predominantly on T cells and NK cells. CD58 enables adhesion and costimulation of cytotoxic T cells and is required for anti-tumor immunity. This protein is, however, missing in rodents (Zhang *et al*, 2021). Since CMTM6-deficient human tumor cells have reduced CD58, they can escape anti-tumor immunity, and increased CMTM6 levels are associated with favorable immune checkpoint therapies in some cancers (Miao *et al*., 2023). Altogether, our results indicate that CMTM6 functions differently in human and mouse cells, as it regulates distinct sets of associated proteins beyond PD-L1. Therefore, the therapeutic potential of targeting human CMTM6 must be further evaluated, as mouse tumor models cannot fully recapitulate the function of human CMTM6 in anti-tumor immunity.

## Methods

### Antibodies

Primary antibodies against β-actin (catalogue number #3700), CMTM6 (#90329), Flag (#14793), cleaved caspase-8 (Asp387) (#9429), cleaved caspase-3 (Asp175) (#9664), and caspase-8 (#4790) were purchased from Cell Signaling Technology. Antibodies against CMTM4 (HPA014704) and Flag (M2) (F3165) were from Sigma-Aldrich. Antibodies against murine FAS (#ab271016) and human FAS (#ab133619) were from Abcam. Secondary antibodies, donkey anti-rabbit IgG (H+L) HRP (#711-035-152), goat anti-mouse IgG1 HRP (#115-035-205), and goat anti-mouse IgG2b HRP (#115-035-207), were from Jackson Immunoresearch. Fluorescently labeled antibodies for flow cytometry against CD3 FITC (#100203), CD4 AF700 (#100536), CD8a BV421 (#100738), CD44 PE (#103007), CD62L BV510 (#104441), CD49d AF647 (#103613), murine FAS APC (#152604), human FAS APC (#305611), murine PD-L1 APC (#124312) and human CD58 APC (#330917) were purchased from Biolegend. Antibody against FoxP3 PE-Cy7 (#25-5773-80) was from Thermo Fisher Scientific.

### Cell lines and animal models

ST2 cells were kindly provided by J. Balounova, and HeLa and HEK293T, Phoenix-Eco, and Phoenix-Ampho cells were kindly provided by T. Brdicka (both from the Institute of Molecular Genetics, Prague, Czech Republic). MEFs were isolated from E11.5 mouse embryos and immortalized via lentiviral transduction with the SV40 large T antigen. All cell lines were cultured at 37 °C in a humidified atmosphere with 5% CO2 in complete Dulbecco’s Modified Eagle Medium (DMEM), supplemented with 10% fetal bovine serum (FBS) (Gibco) and penicillin/streptomycin antibiotics (Biosera). Cell lines were regularly tested for Mycoplasma contamination using the Mycoplasmacheck service (Eurofins Genomics).

We thank the Wellcome Trust Sanger Institute for providing the mutant mouse line carrying the Cmtm6^tm1a(EUCOMM)Wtsi^ allele, and INFRAFRONTIER/EMMA (www.infrafrontier.eu, PMID: 25414328) partner, the National Centre for Biotechnology CNB-CSIC, from which frozen mouse sperm was obtained (RRID: IMSR_EM:06094). Associated primary phenotypic information can be found at www.mousephenotype.org. Frozen sperm was used for in vitro fertilization. Mice carrying the targeted *Cmtm6* allele were crossed to the Flp-deleter mouse strain (RRID: IMSR_JAX:005703) from The Jackson Laboratory. The resulting mice, with exon 2 flanked by LoxP sites, were then crossed with the Cre-deleter strain (RRID: IMSR_JAX:007676) from The Jackson Laboratory to obtain mice carrying the *Cmtm6* KO allele. CMTM6^+/+^ and CMTM6^−/−^ male and female littermates used in experiments were generated by breeding heterozygous animals. Mice were housed under specific pathogen-free conditions with a 12/12-hour light/dark cycle, a temperature of 22 ± 1 °C, and a relative humidity of 55 ± 5 %. Genotyping was performed using PCR with Taq polymerase (Top-Bio) and the primer pairs for *CMTM6* wild-type allele (5’-GCTGCTGTTTCTCATTGCTG-3’, 5’-TGTGTCAAACGCTAAGACTCAGA-3’) and KO allele (5’-GCTGCTGTTTCTCATTGCTG-3’, 5’-GCTATGAACTGATGGCGAGC-3’). Animal protocols were approved by the Czech Academy of Sciences, Czech Republic.

### Generation of KO cell lines

KO cell lines were generated using the CRISPR-Cas9 approach. Single-guided RNAs (sgRNA) were designed using a CHOPCHOP web tool (Labun *et al*, 2019). The following sgRNA sequences were used: mouse CMTM4 (5’-GAAGTAGAGGCCTTCGCACG-3’), human CMTM6 series#1 (5’-GGTGTACAGCCCCACTACGG-3’), human CMTM6 series#2 (5’-GTGAGAACGCGCCGGAGCAAT-3’), mouse FAS (5’-GGCATGGTTGACAGCAAAAT-3’), human FAS series#1 (5’-GATCCAGATCTAACTTGGGG-3’), human FAS series#2 (5’-GACTGCGTGCCCTGCCAAGAA-3’). The sgRNA sequences were cloned into pSpCas9(BB)-2A-GFP (PX458) vector, kindly provided by Feng Zhang (Addgene plasmid #48138) (Ran *et al*, 2013), and transfected into cell lines using Lipofectamine 2000 (Thermo Fisher Scientific). GFP-positive cells were isolated using a FACSAria III cell sorter (BD Bioscience) and subcloned. Target protein expression was assessed by immunoblotting.

### DNA cloning and viral transduction

Coding sequences of various proteins were synthesized using the GeneArt Gene Synthesis service (Thermo Fisher Scientific) and cloned into the retroviral pBabe vector expressing GFP selection marker under the SV40 promoter (kindly provided by M. Hrdinka, University Hospital Ostrava, Czech Republic). Transmembrane and juxtamembrane domains of mouse FASL (AA165-193) and mouse TRAILR2 (AA175-209) were swapped to map interaction with CMTM6. All constructs were sequenced.

Phoenix-Eco cells (for mouse cell transduction) or Phoenix-Ampho cells (for human cell transduction) were transfected with the retroviral vectors using Lipofectamine 2000 (Thermo Fisher Scientific) to produce retroviral particles. Virus-containing supernatants were collected, filtered through a 0.2 μm filter, and used to transduce target mouse (ST2, MEFs) or human (HeLa) cell lines in the presence of polybrene (6 μg/ml, Sigma-Aldrich). Transduction was facilitated by spinfection (2,500 rpm, 45 min, 30 °C). Successfully transduced cells were sorted for GFP expression using a FACSAria IIu cell sorter (BD Biosciences).

### Tandem affinity purification and mass spectrometry analysis

For each experimental condition, murine ST2 or human HeLa cells were cultured on six 15-cm dishes. Cells were washed with serum-free DMEM and lysed on ice for 30 min in lysis buffer (30 mM Tris, pH 7.4, 120 mM NaCl, 2 mM KCl, 2 mM EDTA, 10% glycerol, 10 mM chloroacetamide, Complete protease inhibitor cocktail, and PhosSTOP tablets (Roche)) containing 1% n-Dodecyl-β-D-Maltoside (DDM) (ThermoFisher Scientific). Lysates were cleared by centrifugation (21,130 g, 30 min, 2 °C) followed by a two-step immunoprecipitation.

In the first step, lysates were incubated overnight with 50 μl of anti-Flag M2 affinity agarose gel (Sigma). The next day, the beads were washed three times with lysis buffer containing 0.1% DDM. Bound proteins were eluted overnight in 250 μl of lysis buffer containing 1% DDM and 100 μg/ml 3xFlag peptide (Sigma). A second elution was performed for an additional 6 hours, and both eluates were pooled. The second purification step was overnight incubation of the pooled supernatants with 50 μl of Strep-Tactin sepharose beads (IBA Lifesciences). Samples were subsequently washed three times with lysis buffer containing 0.1% DDM and once with lysis buffer alone. Finally, bound proteins were eluted by incubating the beads with 220 μl of elution buffer (2% sodium deoxycholate, 50 mM Tris, pH 8.5). The eluted protein samples were reduced by adding 10 mM Tris(2-carboxyethyl)phosphine and 40 mM chloroacetamide and heated (95 °C, 10 min).

Samples were further processed using SP3 beads (Hughes *et al*, 2019). SP3 beads were added to a mixture of proteins in the lysis buffer, and protein binding was induced by the addition of ethanol to a 60 % (vol./vol.) final concentration. Samples were mixed and incubated for 5 min at RT. After binding, the tubes were placed into a magnetic rack, and the unbound supernatant was discarded. Beads were subsequently washed two times with 180 μl of 80% ethanol. After washing, samples were digested with trypsin in 100 mM Triethylammonium bicarbonate (37°C, overnight). After digestion, samples were acidified with 1% trifluoroacetic acid, and peptides were desalted using in-house-made stage tips packed with C18 disks (Empore) (Rappsilber *et al*, 2007).

LC/MS analysis was performed using a Nano Reversed phase column (Ion Opticks Aurora Ultimate XT 25 cm x 75 μm ID, C18 UHPLC column, 1.7 μm particles, 120 Åpore size). Mobile phase buffer A was composed of 0.1% formic acid in water. Mobile phase B was composed of 0.1% formic acid in acetonitrile. Samples were loaded onto the trap column (Acclaim PepMap300, C18, 5 μm, 300 ÅWide Pore, 300 μm x 5 mm, 5 Cartridges) for 4 min at a flow rate of 15 μl/min. The loading buffer was composed of 2% acetonitrile and 0.1% trifluoroacetic acid in water. Peptides were eluted with Mobile phase B gradient from 4% to 35% for 60 min. Eluted peptide cations were converted to gas-phase ions by electrospray ionization and analysed on a Thermo Orbitrap Ascend. Survey scans of peptide precursors from 350 to 1400 m/z were performed at 120K resolution (at 200 m/z) with a 5 × 10^5^ ion count target. Tandem mass spectrometry (MS) was performed by isolation at 1,5 Th with the quadrupole, HCD fragmentation with normalized collision energy of 30, and rapid scan MS analysis in the ion trap. The MS/MS ion count target was set to 10^4^, and the max injection time was 35 ms. Only those precursors with charge state 2–6 were sampled for MS/MS. The dynamic exclusion duration was set to 45 s with a 10 ppm tolerance around the selected precursor and its isotopes.

MS data were analyzed and quantified with the MaxQuant software (version 2.4.13.0) (Cox *et al*, 2014). The false discovery rate (FDR) was set to 1% for both proteins and peptides with a minimum peptide length of seven amino acids. The Andromeda search engine was used to search MS/MS spectra against the murine or human Swiss-Prot database (downloaded from Uniprot in October 2024). Trypsin specificity was set as C-terminal to Arg and Lys, also allowing the cleavage at proline bonds and a maximum of two missed cleavages. N-terminal protein acetylation, carbamidomethylation, and Met oxidation were included as variable modifications. Label-free quantification was performed using the Top3 algorithm, which uses the average intensity of the three most intense peptides per protein. Data analysis was performed using Perseus 1.6.14.0 software (Tyanova *et al*, 2016). To exclude common contaminants, we compared the list of potential interactors with the CRAPome database (Mellacheruvu *et al*, 2013).

### Production and testing of recombinant hexameric Fas ligand

Recombinant Hex-FASL consisted of CD33 leader, 6xHis tag, 2xStrep-1xFlag tag, oligomerization domain from mouse adiponectin (AA18-111) (Holler *et al*, 2003), and C-terminal part of mouse FASL (AA136-279). DNA sequence encoding this construct was synthesized using a GeneArt Gene Synthesis service (Thermo Fisher Scientific) and cloned into the pcDNA3.1 vector. The resulting construct was transfected into HEK293T cells using polyethylenimine (PEI) transfection. After three days, the culture supernatant was collected and Hex-FASL was purified using a His GraviTrap TALON column (GE Healthcare) equilibrated with purification buffer (50 mM sodium phosphate, pH 7.4, 300 mM NaCl). The column was washed with purification buffer containing 20 mM imidazole, and Hex-FASL was eluted with purification buffer containing 350 mM imidazole. Following elution, imidazole was removed, and the buffer was exchanged using an Amicon Ultra centrifugal filter with a 10 kDa molecular weight cutoff (Merck). Samples were concentrated by centrifugation and washed three times with purification buffer. The concentration of recombinant protein was measured on a Nanodrop 2000 spectrophotometer (Thermo Fisher Scientific). The purified protein solution was mixed with glycerol to a final concentration of 50% and stored at −80 °C for long-term preservation. The purity of the recombinant ligand was assessed by SDS-PAGE followed by staining with Coomassie InstantBlue (Expedeon).

### Cell stimulation, Immunoprecipitation, and immunoblotting

For analysis of FASL-induced caspase cascade, cells were washed with DMEM and stimulated with 500 ng/ml of Hex-FASL as indicated. After stimulation, cells were lysed in lysis buffer containing 1% DDM for 30 min on ice and cleared by centrifugation (21,130 g, 30 min, 2 °C). Cleared lysates were mixed with 4x SDS sample buffer (250 mM Tris pH 6.8, 8% SDS, 40% glycerol, 0.2% bromophenol blue), reduced with 50 mM dithiothreitol (DTT), and heated (92 °C, 3 min). Samples were analyzed via immunoblotting using the indicated antibodies and imaged using the ChemiDoc MP imaging system (Bio-Rad).

For isolation of proteins via immunoprecipitation, cells were washed with DMEM, solubilized in lysis buffer containing 1% DDM on ice for 30 min, and cleared by centrifugation (21,130 g, 30 min, 2 °C).

A portion of the cleared lysates was mixed with 4x SDS sample buffer, reduced with 50 mM DTT, and heated (92 °C, 3 min) to prepare the lysate samples. To isolate exogenously expressed SF-tagged proteins, lysates were subjected to overnight immunoprecipitation using anti-Flag M2 affinity agarose gel (Sigma). To analyze FAS interacting partners, cellular lysates were incubated with either FAS-specific or isotype control antibody for 1 hour at 4 °C, followed by overnight immunoprecipitation with Protein A/G PLUS agarose (Santa Cruz Biotechnology). The next day, agarose beads were washed three times with lysis buffer containing 0.1% DDM, mixed with 4x SDS sample buffer, reduced with 50 mM DTT, and heated (92 °C, 3 min). Samples were analyzed via immunoblotting.

### Analysis of cell death induction

Cells were seeded in 96-well plates in DMEM supplemented with 10% FCS and ATB at a density of 8,000 cells per well. The following day, cells were stimulated or not with recombinant Hex-FASL (500 ng/ml) in the presence of eTox Red Dye (250 nM, Agilent) to visualize dead cells. The cells were monitored every two hours over a 24-hour period using an Incucyte SX1 live-cell analysis system (Sartorius). In each experiment, samples were analyzed in triplicate, and four images were captured per well. Cell death induction was quantified as the ratio of red fluorescent area to total cell confluency.

### Flow cytometry

For the analysis of cell lines, cells were resuspended in FACS buffer (PBS, 2% FBS, 0.1% NaN3), stained using fluorescently labeled antibodies for 30 min on ice. Cells were washed with ice-cold FACS buffer. Surface expression of selected proteins was measured on the Bricyte E6 flow cytometer. For the analysis of immune cell populations, spleens and peripheral lymph nodes were collected post-mortem from 8-12-week-old mice that had been humanely euthanized for analysis. The organs were dissociated into single-cell suspensions using nylon mesh. Splenic cells were incubated in ACK lysis buffer (150 mM NH4Cl, 10 mM KHCO3, 0.1 mM EDTA-Na2, pH 7.4) to remove red blood cells. Cells were resuspended in FACS buffer and stained on ice with the LIVE/DEAD near-IR dye (Life Technologies), followed by fixation with Foxp3 transcription factor staining buffer set (Thermo Fisher Scientific). Fixed samples were stained with the anti-mouse antibody mixture consisting of CD3 FITC, CD4 AF700, CD8a BV421, CD44 PE, CD62L BV510, CD49d AF647, FAS APC, and FoxP3 PE-Cy7. Samples were measured on an Aurora flow cytometer (Cytek).

### Statistics and data analysis

The indicated statistical analysis was performed using Prism (GraphPad Software). The flow cytometry data were analyzed by FlowJo Software (TreeStar). Immunoblotting data were analyzed using Image Lab Software (Bio-Rad).

## Acknowledgment

We thank J. Stefanovic for technical assistance. Mass spectrometry analyses were performed in the OMICS Mass Spectrometry Core Facility, Faculty of Science, Charles University, at the Biocev research center. The project was supported by a Czech Science Foundation grant (21-25251S) awarded to PD. TS and MP were students at the First Faculty of Medicine at Charles University in Prague and supported by a grant from the Ministry of Education, Youth and Sports of the Czech Republic (SVV 260763).

## Author Contributions

TS planned, performed, and analyzed the majority of the experiments. MP contributed to flow cytometry analysis. HK contributed to the analysis of protein interactions. TT contributed to the analysis of cell death. OS provided animal models. PD supervised the study and wrote the manuscript. All authors commented on the manuscript draft.

## Supplementary Figures

**Figure S1.**
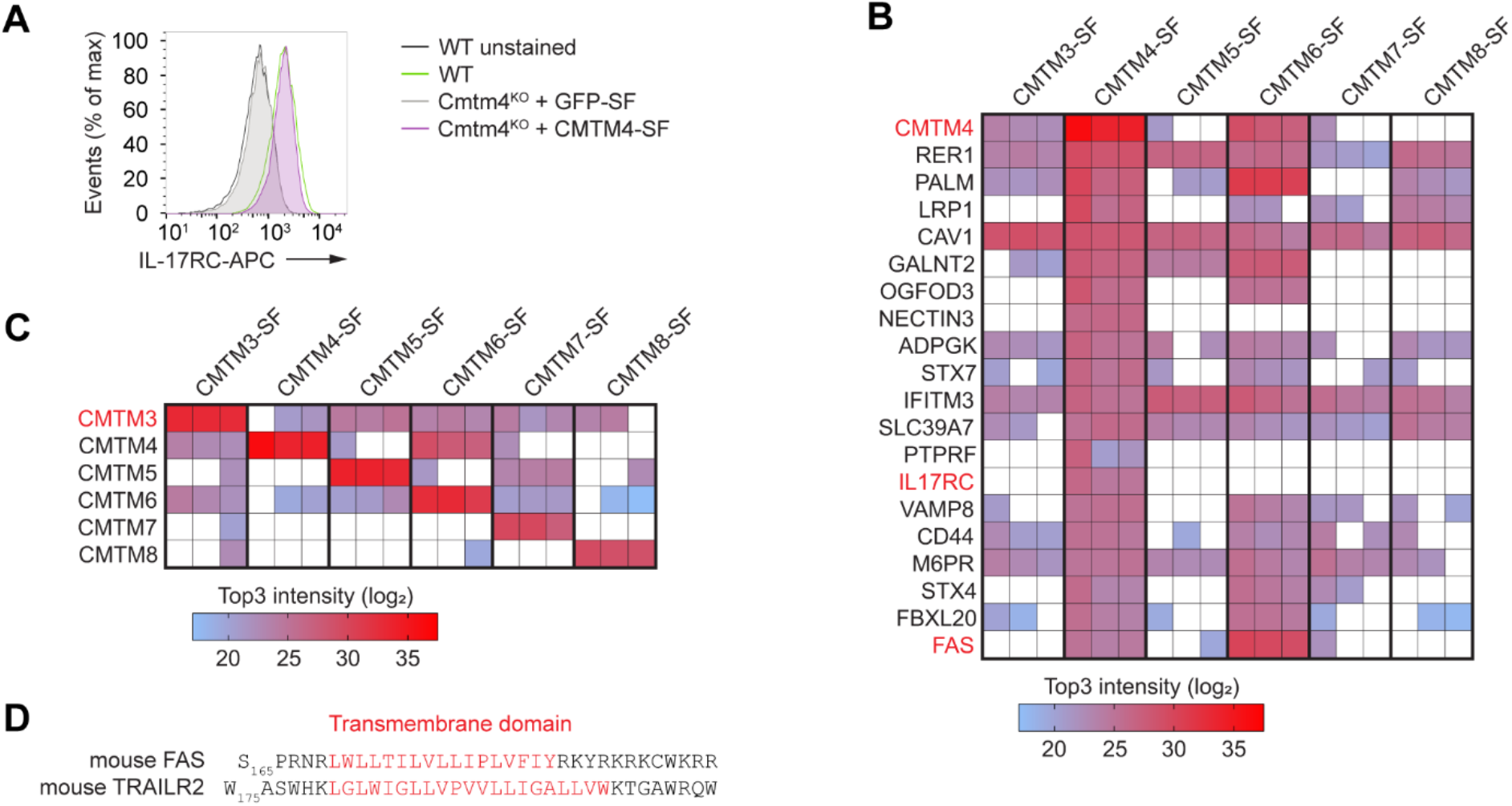
Analysis of the CMTM family interactome. **A**.Flow cytometry analysis of surface IL-17RC expression in WT ST2 cells or *Cmtm4*^KO^ cells transduced with expression vectors encoding GFP-SF or CMTM4-SF. **B**. Mass spectrometry analysis of the indicated Strep-Flag (SF)-tagged CMTM family members isolated from mouse ST2 cells via tandem affinity purification. The heatmap displays the most abundant CMTM4-associated proteins that were not detected in GFP-SF control samples, based on Top3 intensities from 3 independent experiments. **C**. Heatmap showing Top3 intensity values for the associations between various CMTM family members. **D**. Sequence comparison of murine FAS and TRAILR2 transmembrane (in red) and juxtamembrane domains.

**Figure S2.**
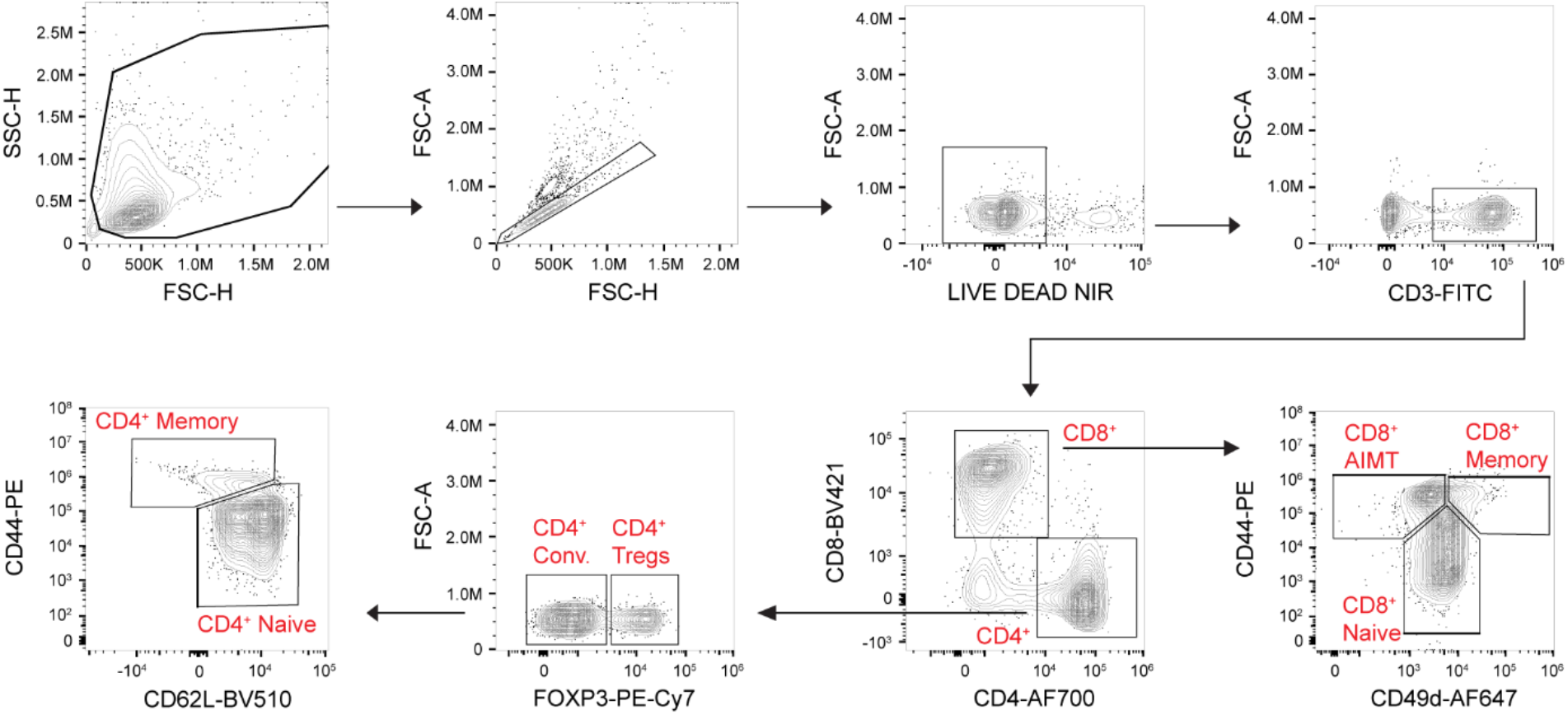
Mouse T cells gating strategy. The gating strategy used for the analysis of T cells isolated from the spleen (Figure 3) and peripheral lymph nodes (Figure S3).

**Figure S3.**
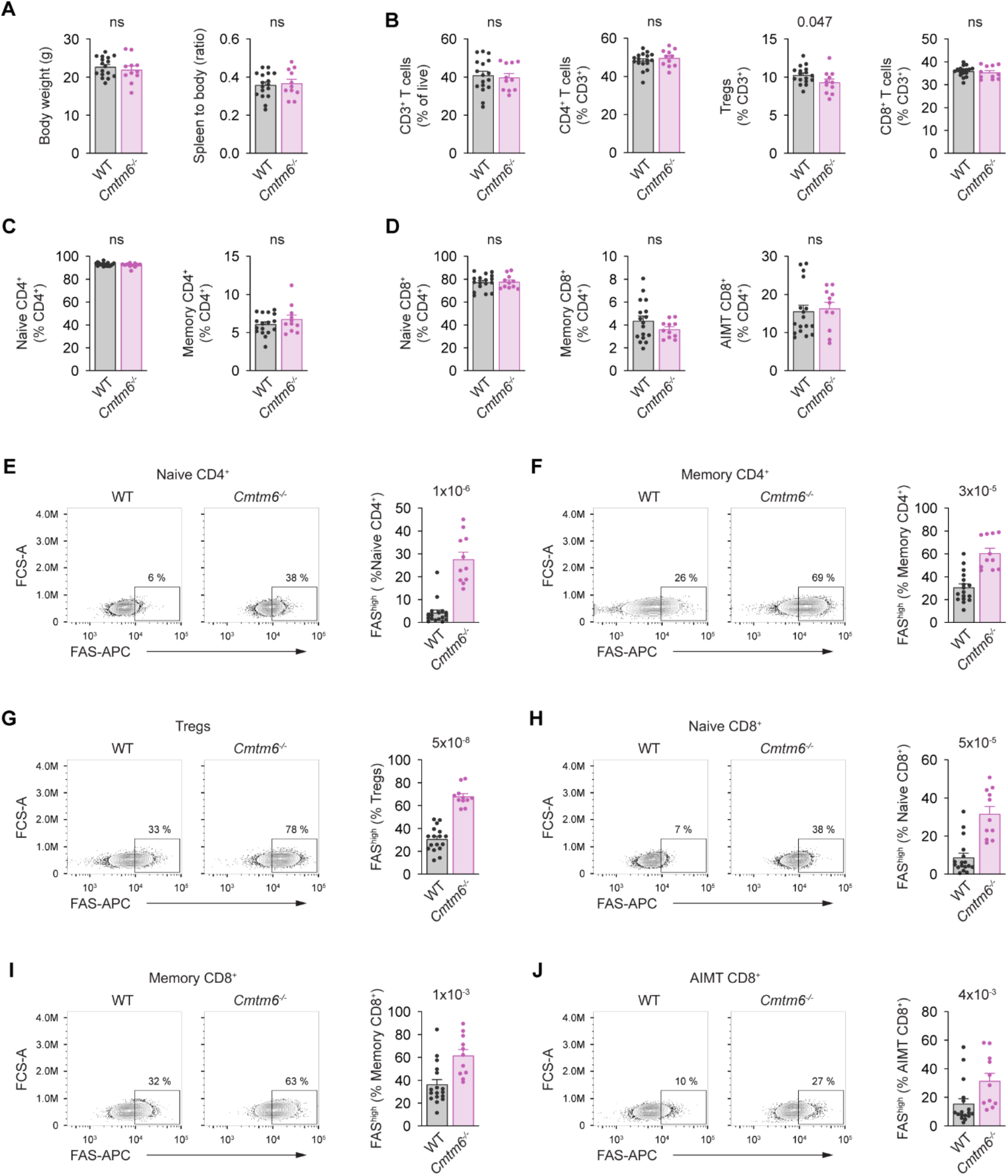
CMTM6 suppresses FAS expression in T cells isolated from mouse lymph nodes. **A**.Body weight and spleen-to-body ratio in 8-12-week-old WT and *Cmtm6*^*-/-*^ mice. **B**. Flow cytometry analysis of T cells isolated from peripheral lymph nodes of 8-12-weeks-old *WT* and *Cmtm6*^*-/-*^ mice. T cells (gated as CD3^+^) were subdivided into conventional CD4^+^ (CD4^+^, FOXP3^−^), Tregs (CD4^+^, FOXP3^+^), and conventional CD8^+^ (CD8^+^). **C**. Conventional CD4^+^ T cells were further gated as naïve (CD4^+^, FOXP3^−^, CD44^−^, CD62L^+^) and memory (CD4^+^, FOXP3^−^, CD44^+^, CD62L^−^). **D**. Conventional CD8^+^ T cells were gated as naïve (CD8^+^, CD44^−^), memory (CD8^+^, CD44^+^, CD49d^+^), and antigen-inexperienced memory T (AIMT) (CD8^+^, CD44^+^, CD49d^−^) cells. **E-J**. FAS expression in the indicated subsets of T cells isolated from peripheral lymph nodes. Data are presented as mean + SEM, n=8 mice per group. Two-tailed Mann-Whitney test. ns, not significant.

**Figure S4.**
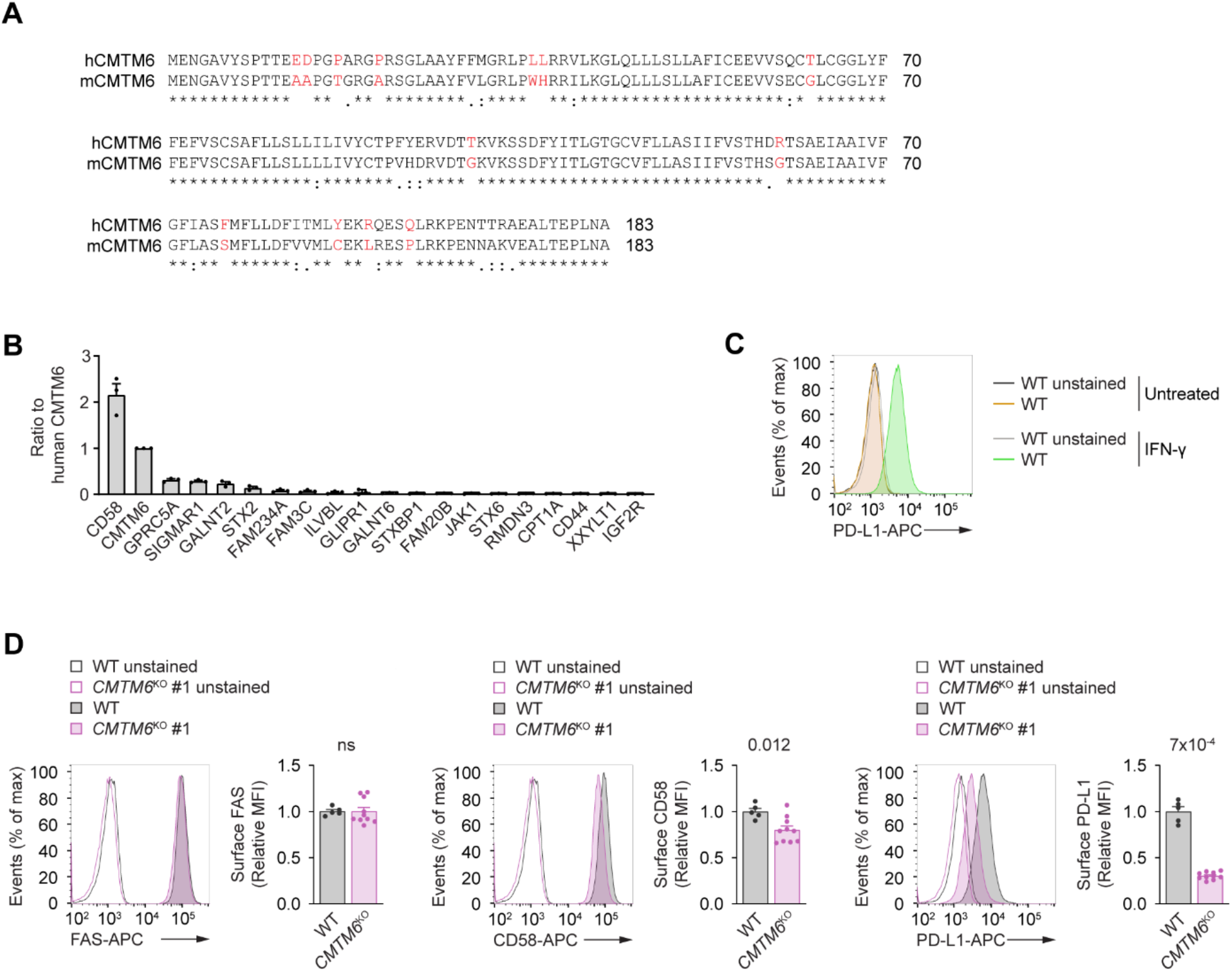
CMTM6 does not regulate human FAS. **A**.Comparison between murine and human CMTM6 sequences. **B**. Mass spectrometry analysis of the SF-tagged human CMTM6 isolated from unstimulated HeLa cells via tandem affinity purification. The 20 most abundant CMTM6 interactors not detected in control GFP-SF samples are shown, based on Top3 intensity from 3 independent experiments. **C**. Flow cytometry analysis of surface PD-L1 expression in WT HeLa cells stimulated or not with human IFN-γ (100 ng/μl). **D**. Flow cytometry analysis of surface FAS, CD58 in untreated and PD-L1 in IFNγ-stimulated WT and *CMTM6*^KO^ HeLa cells. Data are representative of five (D) independent experiments using two different knockout clones. Two-tailed Mann-Whitney test. ns, not significant. MFI, median fluorescent intensity.

